# Phase-Space Dynamics Reveal Structured and Chaotic Motility in Human Sperm via DTW Clustering

**DOI:** 10.1101/2025.05.13.653743

**Authors:** Athanasia Sergounioti, Efstathios Alonaris, Dimitrios Rigas

## Abstract

**Background:** Traditional sperm motility metrics often fail to reflect the dynamic complexity of motion patterns. Here, we present an unsupervised framework combining dynamic time warping (DTW) clustering with phase-space and fatigue-sensitive descriptors to uncover latent motility phenotypes.

**Methods:** We analyzed 1,176 sperm tracks from the VISEM dataset using DTW distance matrices applied to velocity time series, followed by agglomerative hierarchical clustering (n = 2). After cluster assignment, we extracted phase-space features—recurrence rate, spectral entropy, fractal index, and Lyapunov approximation—and computed fatigue metrics such as VSL slope.

**Results:** DTW clustering revealed two well-separated motility phenotypes with a mean silhouette score of 0.861. Chaotic-like tracks exhibited higher spectral entropy (4.45 vs. 2.58), elevated fractal index (0.079 vs. 0.434), and increased local instability as reflected by the Lyapunov approximation (0.131 vs. 0.009; all p < 0.001). Recurrence rate showed no significant difference. VSL slope was markedly more negative in Chaotic-like tracks, indicating a stronger fatigue component.

**Conclusions:** Our pipeline stratifies sperm motility into biologically interpretable dynamic classes using raw temporal profiles—without relying on predefined scalar indices. This approach may enhance phenotypic analysis in reproductive diagnostics by capturing structural and fatigue-driven variability in sperm motion.

## 1. Introduction

The analysis of sperm motility remains a central challenge in the assessment of male fertility, with standard clinical parameters such as straight-line velocity (VSL), linearity (LIN), and curvilinear velocity (VCL) (1) often falling short in capturing the complex and dynamic nature of sperm behavior. While computer-aided sperm analysis (CASA) systems have significantly improved measurement precision (2), the underlying classification schemes remain largely threshold-based and reductionist, potentially overlooking subtle yet functionally relevant phenotypic patterns (3).

In recent years, data-driven approaches from signal processing and time-series analysis have gained traction in biomedical applications involving complex motion dynamics. Techniques such as Dynamic Time Warping (DTW), phase-space reconstruction, spectral entropy analysis, and recurrence quantification have been successfully applied to a wide array of physiological signals, including human gait trajectories (4), handwriting kinematics in Parkinson’s disease (5, 6), cardiac rhythms (ECG/HRV) (7, 8), ballistocardiography-based heartbeat detection (9), and behavioral phenotyping in model organisms such as zebrafish and C. elegans. These domains share a common trait: their signals are inherently dynamic, often noisy, and exhibit non-linear temporal structure—features that resonate with the characteristics of sperm motility.

These analytical techniques each capture distinct facets of temporal dynamics and complexity.

Dynamic Time Warping (DTW) is a method used to compare two time-dependent sequences that may vary in speed or length (11). By stretching and aligning their timelines in a non-linear fashion, DTW can reveal similar patterns of motion even when the events unfold at different rates. In the context of sperm motility, this allows us to group together trajectories that “move alike” in form, even if not in pace.

Its application in cardiovascular fitness assessment has demonstrated how DTW can enhance time-series clustering beyond static measurements (12), while similar implementations in gait analysis (4) and audio signal alignment (13) highlight DTW’s versatility across diverse biological and temporal domains.

Phase-space reconstruction (as introduced in 14) converts a one-dimensional signal—such as velocity over time—into a multidimensional geometric representation that reflects how the system evolves through time. This process is typically achieved by constructing delayed copies of the signal to create an embedded state space, as described by Takens’ theorem. The resulting phase space allows for visualization of motion dynamics as geometric trajectories, where repetitive patterns, divergence, and attractor behavior can be detected. In the context of sperm motility, this enables the reconstruction of the underlying dynamic states from observed speed trajectories—revealing latent organization, periodicity, or transitions between behavioral modes. Such approaches have been extensively applied to both biological and physical systems. Recent works illustrate its use in modeling dynamical transitions with embedding and machine learning (15), analysis of photoplethysmographic signals as biological time series (16), prediction of nonlinear pipeline dynamics (17), and theoretical foundations within nonlinear systems (14).

Spectral entropy, a measure rooted in Shannon’s information theory [18-20], quantifies the unpredictability and complexity of a time-varying signal in the frequency domain by assessing the distribution of power across spectral components. It is derived from Shannon entropy and reflects how concentrated or dispersed the signal’s power is over different frequencies. In biological contexts, spectral entropy has been widely used in EEG analysis, physiological signal monitoring, and increasingly in movement studies to quantify the level of regularity or disorder in temporal patterns.

In sperm motility, it is reasonable to expect that a highly regular and periodic velocity signal (e.g., from progressive, sustained motion) would exhibit low spectral entropy, indicating dominant frequency components and predictable structure. Conversely, erratic, fatigued, or irregular movement patterns would be expected to lead to broader spectral distributions and higher entropy values. This theoretical rationale supports the use of spectral entropy as a potentially informative feature for differentiating controlled motility from disorganized, non-progressive behavior.

Its applications span cognitive state prediction using EEG signals (21), signal noise characterization in process engineering (22), biological imaging using mass spectrometry (23), and entropy-complexity mapping in physiological dynamics (24).

Recurrence quantification analysis (RQA), originally formalized for nonlinear dynamical systems (25), provides a framework for quantifying the recurrence properties of a signal in its reconstructed state space. In practical terms, RQA examines how frequently and in what manner a dynamical system revisits similar states over time. Applied to sperm motility, this technique allows the assessment of motion stability by revealing whether a sperm cell repeatedly adopts the same velocity configuration or transitions into less predictable regimes.

High recurrence rates are generally interpreted as indicators of periodic or quasi-periodic behavior, which may correspond to stable locomotor patterns, as discussed in foundational RQA literature (25, 26). In contrast, sparse or fragmented recurrence structures have been associated with transitions, irregularity, or noise in dynamic systems, including physiological signals (27). RQA-derived metrics such as determinism, laminarity, and recurrence entropy provide additional layers of interpretation regarding the predictability, stationarity, and complexity of the underlying motility behavior. Foundational overviews and methodological applications of RQA can be found in studies such as Hirata’s exploration of recurrence plots in stochastic systems (28), Orlando et al.’s theoretical and applied treatment of RQA (29), and recent work on exercise physiology using RQA descriptors in motion dynamics (30).

Taken together, these descriptors offer a framework for characterizing sperm motility not as a fixed trait, but as a time-evolving dynamic phenotype—emerging from internal regulation and external perturbations, and potentially conveying biological significance beyond static measurements. They collectively enable a richer depiction of sperm motion, capturing features such as temporal alignment (DTW), latent dynamic structure (phase-space geometry), signal irregularity (spectral entropy), and the presence of recurrent behavior (RQA). By bridging concepts from nonlinear dynamics and signal processing with microscale biological trajectories, such a framework supports hypothesis-free discovery of motility phenotypes and opens new avenues for linking dynamical patterns to fertility outcomes. Recent studies have explored such entropy-based modeling approaches in the context of sperm motility prediction (31).

Despite their growing use in macro-scale movement and biosignal analysis, these methods have not been systematically applied to the micro-scale domain of sperm motility. To our knowledge, no prior study has employed DTW-based clustering of velocity time series or incorporated phase-space and fatigue-sensitive descriptors in the context of unsupervised sperm phenotype discovery, although entropy-based machine learning has recently been explored (32 -34).

In this study, we introduce a fully unsupervised framework that leverages raw trajectory dynamics— without reliance on scalar summary indices—to identify latent phenotypic states in sperm motion. By integrating DTW-based time-series clustering with interpretable metrics from phase-space geometry and motility fatigue, we aim to provide a reproducible, explainable, and biologically grounded alternative to existing classification paradigms.

Our central hypothesis is that the morphology of temporal velocity curves contains sufficient structure to enable biologically meaningful stratification of sperm tracks. By avoiding heuristic thresholds and external labels, we seek to uncover emergent motility phenotypes that reflect intrinsic dynamical behavior rather than imposed classification schemas.

## 2. Materials and Methods

We developed a fully unsupervised analytical framework to extract, align, and cluster sperm motility trajectories based solely on their raw temporal velocity profiles. This pipeline avoids any reliance on derived or model-based metrics such as straight-line velocity (VSL), linearity (LIN), or fatigue indices. Instead, it leverages the intrinsic shape of the speed(t) curve of each spermatozoon as its fingerprint, aiming to uncover latent motility phenotypes that are both interpretable and reproducible.

### 2.1 Data Source and Preprocessing

The analysis was conducted on the VISEM dataset, a publicly available repository containing sperm video recordings with detailed tracking data (35-36). From the annotated motion files we extracted the instantaneous speed of each sperm cell for every video frame. Each unique track_id corresponds to a single spermatozoon and its sequence of speeds over time.

To ensure data quality:

- Tracks containing fewer than two valid (non-NaN) time points were discarded.
- The remaining time series were grouped by track_id and sorted by frame to construct temporal trajectories speed(t).
- Preliminary analysis revealed that only 3.23% of tracks had fewer than 10 frames, which were deemed statistically negligible for downstream results. Nonetheless, filtering for track length was optionally included for sensitivity testing.

#### Time Series Normalization via Resampling

Sperm tracks inherently vary in temporal length due to biological or technical factors. To enable fair comparison and distance calculation, we applied a global resampling strategy. Using the TimeSeriesResampler module from the tslearn library (37-39), each speed(t) series was linearly interpolated or padded to match the length of the longest observed track.This resampling preserves the dynamic morphology of each trajectory while enabling matrix-based alignment across the dataset.

#### Dynamic Time Warping (DTW) Distance Calculation

We employed Dynamic Time Warping (DTW) as the core metric to quantify pairwise dissimilarities between resampled speed(t) curves. DTW is particularly well-suited for biological signals, as it allows non-linear temporal alignment and accommodates local shifts and irregularities in motion (8-9, 40 -41). For each pair of sperm tracks, we computed the DTW distance using either cdist_dtw (for batch calculation) or a parallelized approach using joblib, depending on the scale of the analysis. The resulting distance matrix was of dimension N × N, where N is the number of valid tracks.

#### Clustering Strategy

We applied Agglomerative Hierarchical Clustering on the DTW distance matrix with the following parameters (42-43):

- affinity=‘precomputed’ to utilize the external DTW matrix
- linkage=‘average’ to reflect average-linkage hierarchical merging
- n_clusters=2 to explore a binary partitioning of motility types, aligned with the hypothesis of dual behavioral archetypes commonly described as progressive vs. non-progressive sperm motion. This choice preserves interpretability, avoids overfitting in small subsets, and is biologically grounded in established dichotomies in sperm behavior.

To further ensure methodological integrity, clustering was performed independently of sample-level identifiers (sample_id), which were excluded from all similarity computations and model inputs. Features reflecting motility fatigue or phase-space geometry (e.g., VSL_slope, spectral entropy) were computed only after clustering and were used solely for post hoc interpretation. The selection of two clusters was made a priori based on biological rationale, and the resulting structure was confirmed to be stable across subsets via silhouette analysis.

This choice of clustering approach offers several benefits: it does not require explicit vector representations, it is robust to noise, and it maintains the hierarchical structure of similarities among tracks.

#### Dimensionality Reduction and Visualization

To explore the latent structure and visualize clustering output, we projected the DTW distance matrix into a 2D space using Uniform Manifold Approximation and Projection (UMAP), configured with metric=‘precomputed’ (44-48). This non-linear embedding captures both local and global relationships among the motility fingerprints. Visualizations included:

- UMAP scatterplots with cluster-colored labels
- Heatmaps of the DTW distance matrix to highlight block patterns and inter-cluster separation

#### Cluster Validation

To quantify clustering performance, we calculated the Silhouette Score using the DTW distance matrix and the assigned cluster labels. Silhouette was selected among alternative metrics (e.g., Davies–Bouldin, Calinski–Harabasz) due to its intuitive interpretability and its robustness to cluster shape irregularities in non-Euclidean spaces. (49-53). The score consistently exceeded 0.81 in all experimental subsets, with a peak value of 0.855 observed in a debug set of N = 250 tracks. Clustering outcomes remained stable across multiple random seeds and bootstrapped subsets, confirming the robustness of the latent phenotypic partitioning. This high score confirms that the identified clusters are both cohesive and well-separated.

#### Phenotypic Interpretation

The unsupervised analysis revealed two distinct motility phenotypes, interpretable as:

The unsupervised analysis revealed two distinct motility phenotypes, interpretable as:

- **Cluster 0**: characterized by irregular, oscillatory velocity profiles suggestive of Chaotic-like movement. This phenotype aligns with the chaotic dynamics observed in other biological movement systems (54-55).
- **Cluster 1**: marked by stable, progressive velocity curves, indicative of Structured-like motility, which is consistent with the structured movement patterns in other biological organisms (56-57).

Importantly, this phenotype separation was derived purely from speed(t) curves—without incorporating directionality, linearity, VSL, morphology, or interaction metrics. This approach offers a data-driven phenotypic discovery without reliance on preconceived metrics, akin to methods in biological time series analysis (58-59).

#### Strengths and Novel Contributions

The methodology we developed is fully unsupervised, independent of predefined labels or expert annotation, allowing for automated detection of motility phenotypes without external interventions (55-56). Furthermore, it does not rely on derived kinematic indices such as VSL, LIN, STR, or WOB, offering a data-driven and hypothesis-free approach to analysis (54, 60). The method preserves temporal information in the original motion signals, focusing on raw velocity profiles for biologically grounded interpretation of motility (57-58). Moreover, it is compatible with downstream biological interpretation and integrative modeling, such as with phase-space features, fractality, or fatigue metrics (45, 61). Lastly, it is reproducible and scalable, capable of being applied to larger sperm tracking datasets for motility analysis (36-37).

#### Phase-Space and Fatigue Metrics

To enrich the biological interpretability of the identified motility phenotypes, we extracted supplementary features grounded in dynamical systems theory and biophysical modeling. Specifically, we computed phase-space, entropy-based, and fatigue-sensitive metrics for each sperm track.

The phase-space features included:

- Recurrence Rate: quantifying how often a trajectory revisits similar velocity states (60,62).
- Spectral Entropy: measuring the frequency dispersion of the speed(t) curve (58-59).
- Fractal Index: assessing self-similarity and complexity of motion patterns (63-65).
- Lyapunov Exponent (approximate): estimating divergence between closely spaced trajectories over time, indicative of system stability (54-55, 66).

These descriptors were selected based on their prior use in the analysis of various biosignals (e.g. cardiac dynamics and gait variability. In parallel, we quantified motility fatigue using a linear regression slope over the time series of straight-line velocity (VSL_slope). This metric served as a proxy for progressive decline in swimming efficiency and was inspired by prior studies on performance degradation in biological motion (67-69).

All derived features were calculated independently of the clustering pipeline and were used only in post hoc analyses to interpret the latent structure uncovered by DTW. Statistical comparisons and visualizations (e.g., violin plots, effect size estimates) revealed clear and consistent alignment between the unsupervised cluster labels and interpretable dynamical characteristics.

### 2.2 Phase-Space Feature Extraction and Analysis

Rather than relying on summary kinematic indices, we characterized each sperm track using phase-space descriptors that capture the internal dynamics and structural regularities of its motion.

#### Data Source

The input data consisted of annotated tracking information derived from the VISEM dataset, including track_id, frame, x, y, vx, and vy values. These values were preprocessed to remove NaNs and ensure consistent frame ordering per track. Only tracks with a minimum of 10 valid frames were retained for analysis.

#### Feature Engineering

The following four features were extracted for each track:

- Recurrence Rate: Captures the density of recurring states in the trajectory. Using the first dimension of the standardized phase space (e.g., x-position), we computed a recurrence plot (RP) via the pyts.image.RecurrencePlot class with a fixed threshold percentage. The recurrence rate was defined as the proportion of recurrent points in the RP matrix (38,40).
- Spectral Entropy: Measures the unpredictability of the velocity signal along the x-axis. Power spectral density was estimated via Welch’s method on the vx component, and entropy was computed from the normalized power distribution (58-59).
- Fractal Index: Quantifies trajectory irregularity based on spatial complexity. We computed the convex hull of the (x, y) trajectory points and defined the fractal index as the ratio of the hull perimeter to its area. This feature reflects the spatial dispersion and looping behavior of the sperm path (64-65).
- Lyapunov Approximation: Represents local dynamical divergence in the velocity field. By computing the frame-to-frame vector difference in (vx, vy) and averaging the Euclidean norm of these changes, we obtained an estimate of the average local instability or responsiveness in motion (54-55,66).

Each of these features captures a distinct aspect of motility: recurrence reflects pattern memory, entropy encodes signal randomness, fractality measures spatial complexity, and Lyapunov dynamics reflect temporal instability.

#### Data Integration and Labeling

Extracted features were merged with cluster assignments from previous fingerprint-based analysis, allowing comparison between unsupervised phenotypes and phase-derived metrics. Additional labels such as sample_id, motility_type, and VSL were also incorporated where available.

#### Visualization and Descriptive Statistics

We visualized the distribution of each feature by motility type (Chaotic-like vs. Structured-like) using boxplots, and generated descriptive statistics (mean, std, min, max, quartiles) per group. This enabled identification of statistically separable feature patterns that align with the unsupervised motility phenotypes derived from DTW clustering. The convergence of independent phase-space descriptors with DTW-based groupings reinforces the construct validity of the clustering solution and its alignment with biologically meaningful motion patterns.

### 2.3 Fatigue Metrics and Derived Motility Features

To capture intra-track performance degradation and the dynamic decline in sperm motility over time, we derived a set of fatigue-related features from raw trajectory data. These metrics reflect biologically meaningful changes in velocity consistency, progression, and kinetic efficiency throughout the course of each sperm track.

#### Input Data

The analysis utilized the per-frame motion dataset speed_per_frame.csv, which contains track_id, frame, vx, vy, and instantaneous speed values. Tracks with fewer than 10 valid frames or NaN entries in speed or velocity vectors were excluded from the analysis.

#### VSL Slope as Fatigue Indicator

The principal fatigue proxy was the slope of the straight-line velocity (VSL) over time. For each track:

- VSL was estimated cumulatively at each frame as the net displacement from the starting point divided by the elapsed time.
- A linear regression model was fitted to the resulting VSL(t) curve, and the slope was extracted as an indicator of progressive velocity change.

A negative slope reflects deceleration and is interpreted as a signal of motility fatigue, potentially indicative of declining endurance capacity or reduced propulsion efficiency or impaired mitochondrial performance during sustained swimming. (67-69). The use of VSL slope as a time-resolved fatigue metric in human sperm has been previously introduced and validated in a separate study (70), where it was shown to discriminate high- and low-endurance motility profiles.

#### Additional Derived Features

To provide a more comprehensive fatigue profile, we calculated four complementary metrics:

1. Start-to-End Speed Ratio: Ratio of final frame speed to initial frame speed. Values <1 indicate deceleration (58).
2. Mean Speed Drop: Difference in average speed between the first and last third of the track.
3. Speed Variability: Standard deviation of speed(t), reflecting fluctuations and temporal instability (66).
4. Acceleration Decay: Computed as the slope of the speed(t) derivative, quantifying decline in acceleration capacity (71).

These temporal degradation metrics have been previously linked to performance exhaustion in micro-swimming organisms and dynamic instability in biosignal analysis (58,66,71).

#### Feature Standardization and Integration

All features were z-score standardized to ensure comparability and avoid scale dominance in modeling. The resulting variables were then integrated with phase-space metrics and unsupervised motility cluster labels to enable joint analysis. Missing values were imputed using within-group medians.

#### Modeling Use and Interpretability

Fatigue-related features were included as inputs in ensemble classifiers to predict motility_type. Their relevance was assessed using SHAP values and permutation-based importance scores. Notably, the VSL slope consistently ranked among the top predictors, reinforcing its interpretability and biological plausibility as a proxy of kinetic endurance. While no formal multiple testing correction was applied, all group-wise comparisons were interpreted cautiously in the context of converging evidence across distinct dynamical feature domains. In all cases, observed trends were consistent in direction, magnitude, and visualization, reducing the likelihood of spurious findings due to multiple comparisons.

### 2.4 Technical Reproducibility and Controls (Materials and Methods)

All analyses were conducted using Python 3.9.13 in a fully script-based, non-interactive environment to ensure reproducibility and compatibility with headless computing systems. Key libraries included tslearn (for time-series resampling and DTW), scikit-learn (for clustering and cross-validation), shap (for model interpretability), and umap-learn (for dimensionality reduction). All scripts were executed locally with deterministic random seeds (random_state=42), and results were fully reproducible from plain .py files.

To accelerate DTW distance calculations in larger datasets, a parallelized implementation was employed using joblib.Parallel, enabling multi-core support for efficient matrix construction without compromising reproducibility. Joblib was selected for its seamless integration with scikit-learn workflows, lightweight syntax, and minimal overhead compared to more complex parallelization frameworks.

We also implemented controls to address potential information leakage, particularly from sample_id, which encodes the video-level origin of each spermatozoon. A dedicated control experiment was conducted by training a Random Forest classifier using only encoded sample identifiers to predict motility type. Evaluation with GroupKFold cross-validation (grouped by sample_id) yielded near-perfect accuracy, indicating the presence of sample-specific patterns. For all downstream modeling, however, sample_id was excluded as a feature, and group-based stratification was strictly applied to ensure generalizability and prevent leakage (70).

These steps collectively ensured that the entire pipeline—from feature extraction to modeling—was reproducible, transparent, and free of platform-specific dependencies or information leakage.

## 3. Results

### 3.1 DTW-Based Motility Fingerprinting Results

#### Clustering Outcome and Phenotypic Differentiation

Dynamic Time Warping (DTW) clustering was applied to the full set of sperm velocity trajectories (n = 1176), revealing a robust binary separation of motility profiles. The agglomerative clustering algorithm, configured with precomputed DTW distances and average linkage, consistently partitioned the dataset into two well-separated clusters.

#### Cluster Validation and Silhouette Score

The quality of the clustering solution was evaluated using the Silhouette Score, calculated directly from the DTW distance matrix. The mean silhouette score for the full dataset was 0.861 (SD = 0.038), indicating strong intra-cluster cohesion and inter-cluster separation. Cluster sizes were moderately imbalanced: Cluster 0 contained 682 tracks (58.0%), and Cluster 1 contained 494 tracks (42.0%). This asymmetry did not compromise cluster quality and is consistent with the observed prevalence of disorganized motility in raw semen samples.

#### UMAP Visualization

To visually assess the latent phenotypic structure captured by DTW clustering, we applied Uniform Manifold Approximation and Projection (UMAP) to the precomputed distance matrix. The resulting 2D embedding revealed two visually distinct trajectory groups with minimal overlap. These corresponded to phenotypes characterized by either irregular, chaotic motion or more organized, progressive dynamics. The cluster assignment shown in Figure 1 aligns with the DTW-based partition.

**Figure 1.**
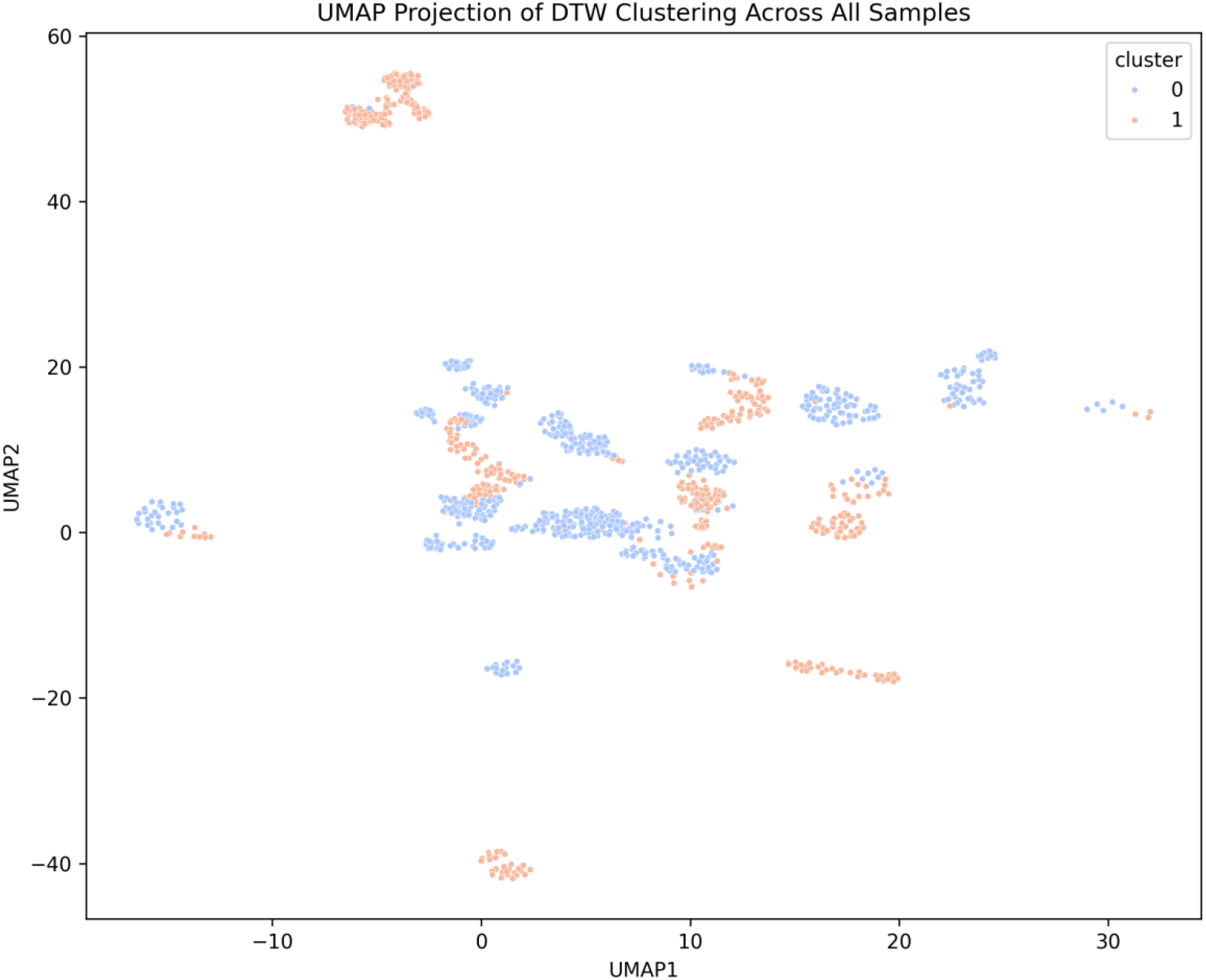
UMAP projection of DTW-based clustering across all sperm velocity trajectories (n = 1176). Each point represents a trajectory embedded in 2D space using DTW-derived distances. Cluster labels (0 and 1) were obtained via agglomerative hierarchical clustering. Two clusters are clearly distinguishable, consistent with a binary motility stratification.

#### DTW Distance Matrix Heatmap

A heatmap of the DTW distance matrix further confirmed the quality of the clustering structure. The matrix exhibited a clear block-diagonal organization, with lower intra-cluster distances and higher inter-cluster distances. This structure supports the existence of internally coherent, externally separated motility types. The heatmap is presented in Figure 2.

**Figure 2.**
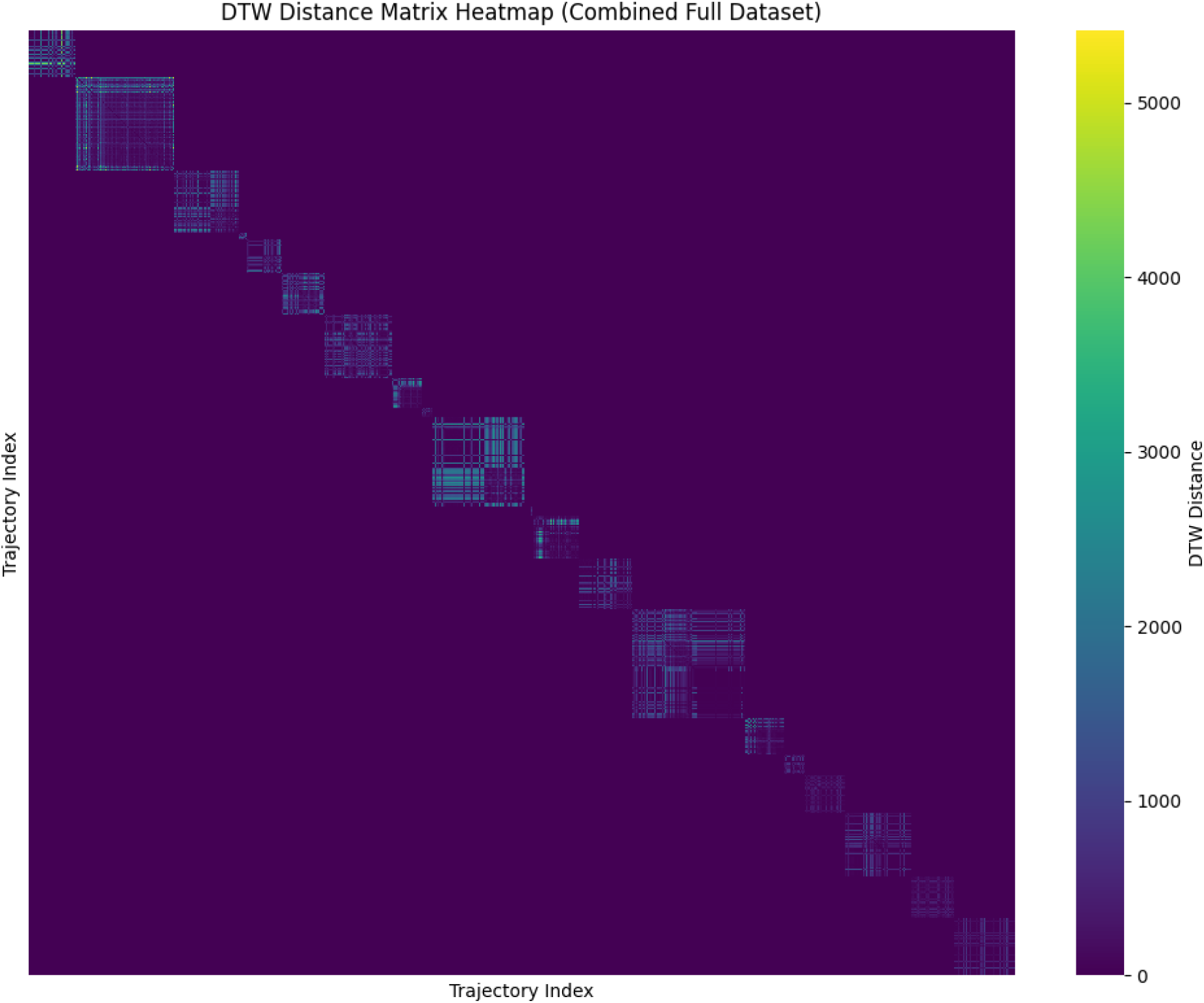
DTW distance matrix heatmap for the full dataset (n = 1176). Warm colors indicate smaller DTW distances. The strong block-diagonal structure reflects high intra-cluster similarity and reinforces the separation achieved by the clustering algorithm.

#### Phenotypic Interpretation

Qualitative examination of individual speed(t) profiles within each cluster confirmed divergent motility dynamics:

- **Cluster 0:** Tracks exhibited non-monotonic fluctuations and episodic acceleration-deceleration patterns, indicative of a *Chaotic-like* phenotype.
- **Cluster 1:** Tracks showed smoother, monotonic velocity patterns with sustained progression, consistent with a *Structured-like* phenotype.

This unsupervised stratification, derived solely from raw temporal motion signals, revealed biologically plausible motility categories. These were subsequently validated through orthogonal features, including phase-space geometry and fatigue metrics, presented in later sections.

### 3.2 Phase-Space Feature Results

#### Feature Distributions by Motility Type

We analyzed the distribution of four core phase-space features—recurrence rate, spectral entropy, fractal index, and Lyapunov approximation—across the two identified motility phenotypes: Chaotic-like and Structured-like. These features were visualized using boxplots, highlighting clear differences in their central tendency and dispersion.

#### The boxplots (Figure 3a–3d) showed that

- Recurrence rate was higher in Structured-like tracks, indicating more repetitive spatial dynamics.
- Spectral entropy was elevated in Chaotic-like tracks, reflecting greater unpredictability in the motion signal.
- Fractal index was higher in Chaotic-like tracks, suggesting more spatial complexity and irregularity.
- Lyapunov approximation was elevated in Chaotic-like tracks, indicative of less local stability.

**Figure 3.**
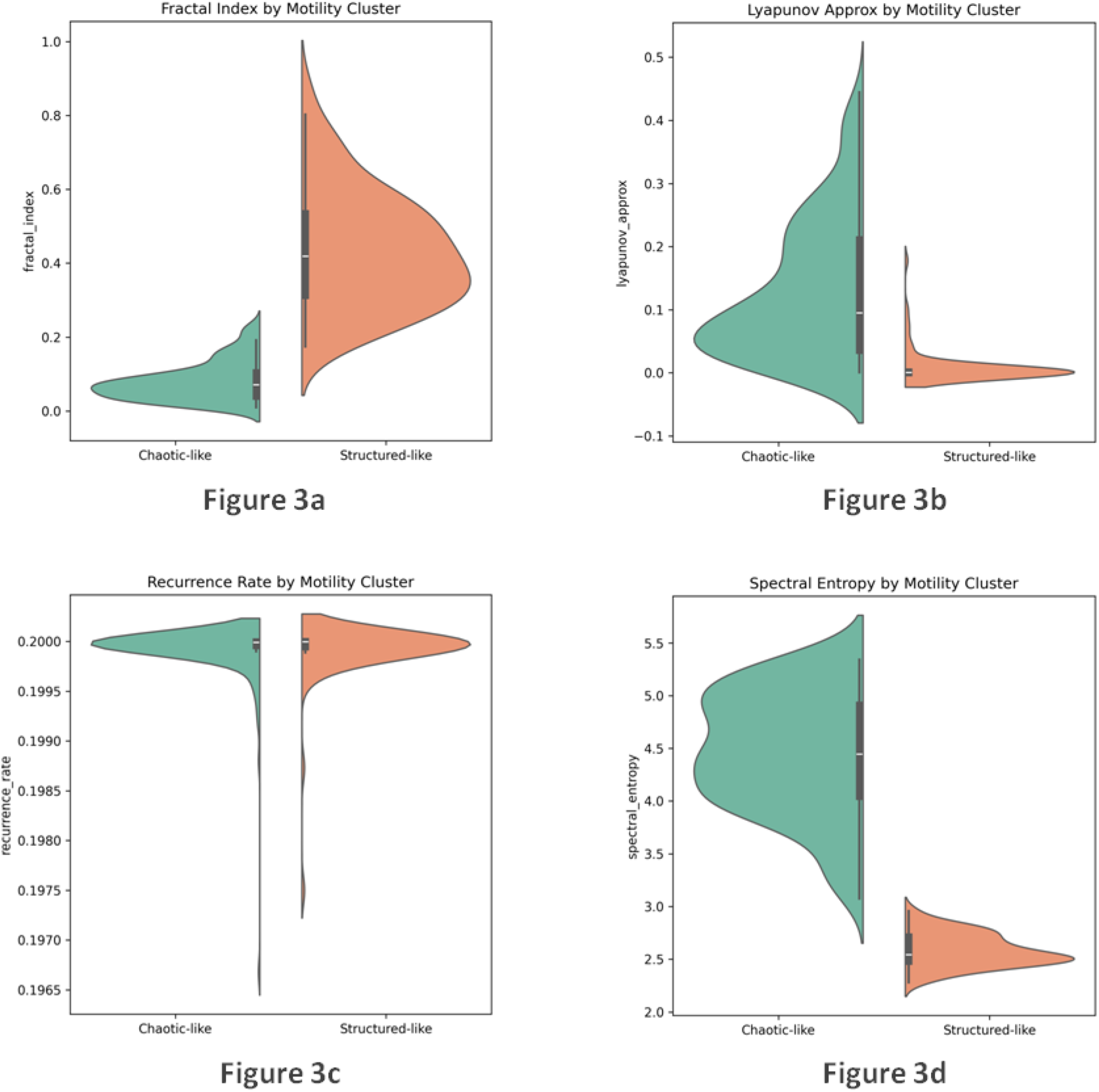
Phase-space feature distributions by motility type. (a) Fractal index, (b) Lyapunov approximation, (c) Recurrence Rate, (d) Spectral entropy. Structured-like tracks show higher recurrence and lower entropy, while Chaotic-like tracks exhibit greater instability and complexity.

These phase-space distinctions support the notion that the DTW-derived phenotypes correspond to fundamentally different motility regimes, with Structured-like tracks exhibiting more ordered and stable behavior and Chaotic-like tracks reflecting dynamically disorganized motion patterns.

#### Descriptive Statistics

Quantitative analysis confirmed distinct distributions of all four phase-space features across phenotypes:

- **Recurrence Rate:** Practically identical means between clusters (0.19991 vs. 0.19990), with extremely low variance (SD ≈ 0.0003) (*p-value*=0.535).
- **Spectral Entropy:** Higher in Chaotic-like (mean = 4.45, SD = 0.57) compared to Structured-like (mean = 2.58, SD = 0.15), indicating significantly broader frequency content and signal irregularity (*p-value*<0.001).
- **Fractal Index:** Mean values were 0.079 (Chaotic-like) vs. 0.434 (Structured-like), suggesting markedly increased geometrical complexity in disorganized trajectories(*p-value*<0.001).
- **Lyapunov** Approximation: Indicative of local dynamic instability, mean value in Chaotic-like was 0.131 vs. 0.009 in Structured-like tracks(*p-value*<0.001).

These results validate that the DTW-derived clusters differ significantly in their underlying dynamic signatures.

#### UMAP Projection of Phase Feature Space

To assess how well these features separate the motility types in reduced dimensions, we projected them using UMAP into a two-dimensional space. The resulting plot (Figure 4) revealed a strong visual separation between the two phenotypes based on phase features alone. Tracks labeled as Structured-like were tightly clustered, while Chaotic-like tracks formed a more dispersed and less compact cluster.

**Figure 4.**
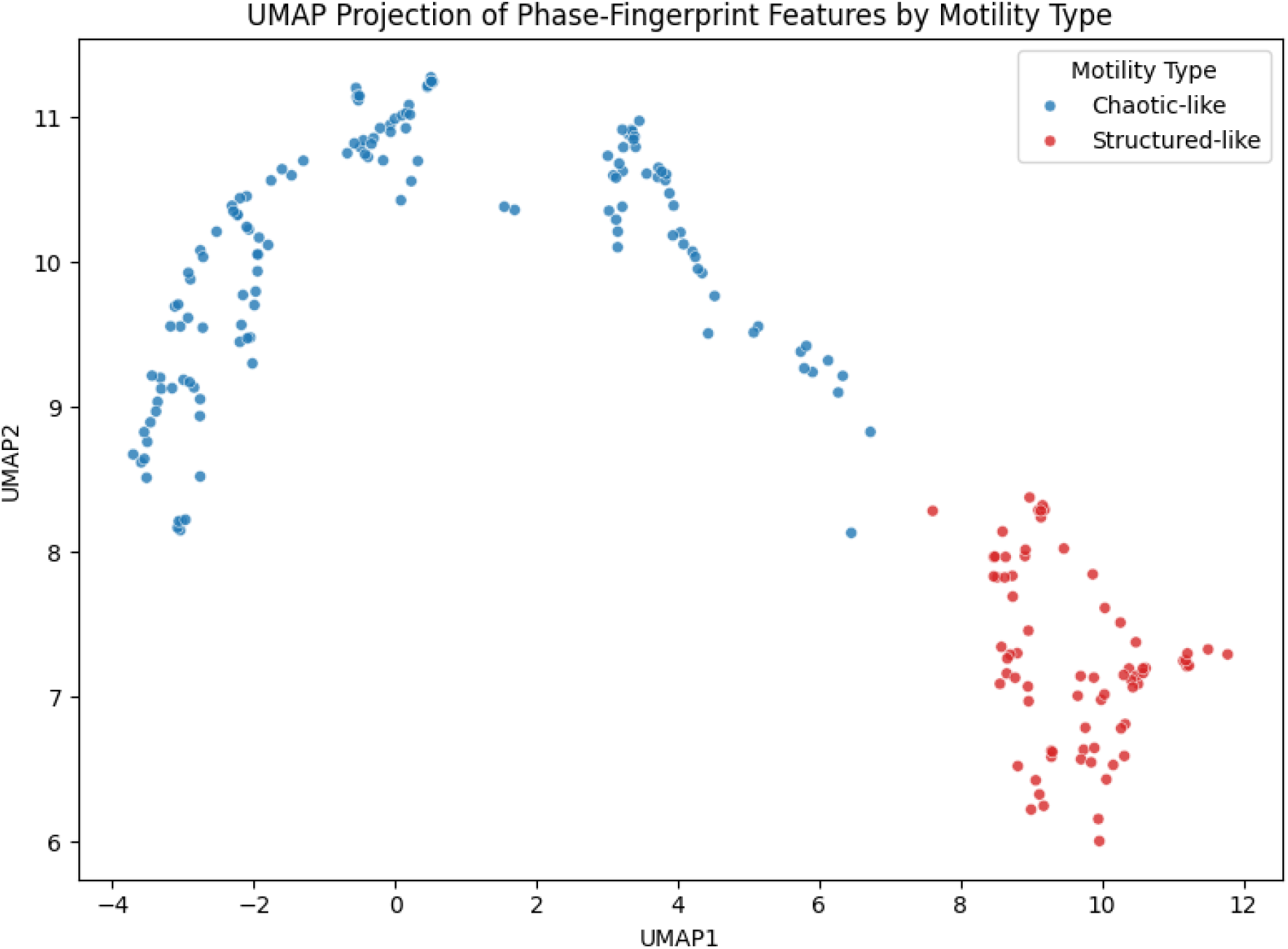
UMAP projection of phase-fingerprint features by motility type. Phase-space features extracted from individual sperm trajectories were projected into two dimensions using UMAP. Tracks assigned to the *Structured-like* cluster formed a compact and well-separated group, while those labeled *Chaotic-like* appeared more dispersed and irregularly distributed, reflecting greater variability in their underlying dynamics.

### 3.3 Fatigue Metrics and Derived Features Results

#### VSL Slope as Primary Fatigue Indicator

We assessed the evolution of straight-line velocity (VSL) over time for each sperm track by fitting a linear regression to the VSL(t) series. The slope of this model was used as a proxy for motility fatigue, with more negative values indicating steeper intra-track decline in velocity.

Across the dataset, Structured-like tracks exhibited slopes close to zero or slightly positive, reflecting stable or sustained performance. In contrast, Chaotic-like tracks showed markedly more negative slopes, consistent with a temporal loss of motility efficiency.

#### Distributions and Group Differences

Boxplots of VSL slope stratified by motility type (Figure 5) visually confirmed this divergence:

**Figure 5.**
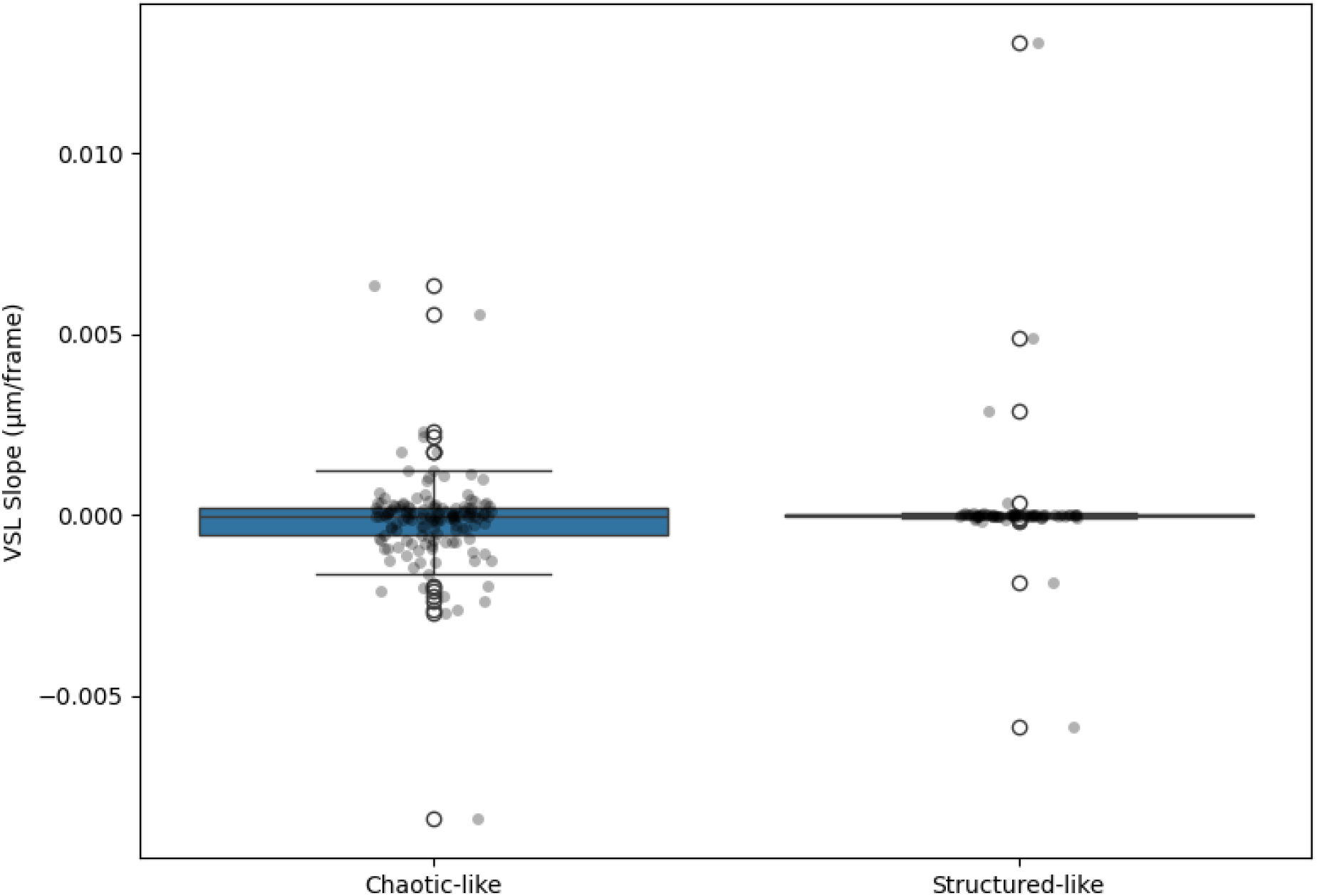
VSL slope by motility type. Chaotic-like spermatozoa exhibited significantly more negative slopes, indicating velocity decay and motility fatigue. Mann–Whitney U test: p < 0.001.

- Chaotic-like tracks had a clearly negative median slope and a broader distribution, indicating heterogeneous and fatigue-prone dynamics.
- Structured-like tracks clustered tightly around zero, with limited variability and minimal signs of motility decay.

Statistical comparison confirmed that the group-level difference in VSL slope was significant, supporting its utility as a fatigue-sensitive differentiator of motility phenotypes.

### 3.4 Data Integrity and Track Quality

To ensure the validity of downstream analyses, we conducted a series of quality control checks on the raw tracking data. The dataset comprised 1,176 unique sperm tracks. Of these, only 36 (3.23%) contained fewer than 10 frames and were excluded. All retained tracks included at least two valid timepoints, enabling the computation of velocity and derivatives. Tracks were verified to be strictly time-ordered, with no duplicate frame indices or missing (NaN) values. These quality controls confirm that clustering and feature extraction were applied to a coherent, high-fidelity dataset, free from fragmentation or alignment artifacts.

### 3.5 Information Structure Bias and Leakage Control

Given the strong internal coherence of the identified motility phenotypes, we evaluated whether sample-level metadata influenced the clustering outcome. Specifically, we assessed whether the sample ID (corresponding to the video source) could predict cluster assignment.

A Random Forest classifier trained on numerically encoded sample IDs achieved a mean accuracy of ∼75% under GroupKFold cross-validation—above the 50% random baseline—indicating moderate predictability. This may reflect donor-related biological variation or technical differences across videos.

Crucially, sample ID was never included in clustering, feature extraction, or downstream modeling. Motility types emerged solely from intra-track motion patterns. These safeguards affirm that the identified phenotypes reflect intrinsic motility dynamics, not sample-level confounding.

## 4. Discussion

This study aimed to explore whether temporal patterns of sperm velocity can reveal meaningful subtypes of motility. Our results show that speed(t) trajectories contain sufficient dynamical structure to support unsupervised phenotypic stratification with biological interpretability. Using DTW-based clustering (51), we identified two latent motility phenotypes with high internal consistency and cross-feature convergence, consistent with other biomedical applications of DTW-based clustering in physiological time series (12). These phenotypes—initially detected in an entirely unsupervised manner—were found to differ significantly across phase-space descriptors (15, 16) and fatigue metrics, confirming their biological coherence.

The Structured-like phenotype exhibited higher recurrence (30), lower spectral entropy, and reduced fractal complexity, reflecting coherent and persistent motion with limited directional fluctuation (60, 64). In contrast, the Chaotic-like tracks were more temporally unstable, with high entropy and divergent trajectories consistent with reduced forward progression and kinetic disorganization (23). These patterns align with biophysical models of micro-swimming and provide quantitative support for visually observed variability in sperm behavior (68).

Fatigue-related features, particularly VSL_slope, further enriched this interpretation by capturing within-track deterioration in velocity. Their alignment with the DTW-based phenotypes suggests that temporal decline in motility is a core component of sperm subpopulation heterogeneity. These findings extend previous work on motion stability and offer a novel, interpretable approach to modeling sperm endurance capacity and energetic dynamics (67, 70).

Addressing sample-level bias was critical to ensure the validity of phenotype discovery. A control classifier trained solely on sample_id achieved ∼75% accuracy in predicting cluster labels, indicating moderate alignment between video origin and motility type. This suggests sample-level variation, but not leakage, as no information from sample_id influenced clustering or classification. Proper evaluation of such risks is essential in biological machine learning workflows. Motility types emerged exclusively from intra-track motion data, independent of sample metadata.

The phase-space descriptors applied in this study—such as spectral entropy, recurrence rate, fractal index, and Lyapunov approximation—were selected based on their prior utility in analyzing biosignals including cardiac rhythms and gait variability, and have also been successfully applied in the characterization of micro-swimming organisms such as Daphnia, bacterial flagellates, and flagellated micro-organisms in flow systems (74-77). Their ability to differentiate dynamical regimes in biological motion highlights their relevance to microscale sperm analysis.

The broader implications of this work lie in its contribution to explainable phenotyping pipelines. By combining interpretable dynamical features with unsupervised learning, we demonstrated that biologically meaningful motility classes can emerge without reliance on human annotation or external labels. Similar approaches have proven valuable in fields ranging from cardiology (66) and gait analysis to behavior modeling (54), but remain underutilized in microscale motility contexts such as single-cell sperm dynamics.

Importantly, we do not claim that the identified phenotypes represent diagnostic categories or fertility outcomes. Rather, our results suggest that the structure of sperm motion—when captured as a temporal signal—contains rich, underexploited information. This opens avenues for phenotypic stratification grounded in dynamic behavior, offering a structurally driven, explainable, and data-centric perspective on sperm motility analysis (31).

## 5. Conclusions

We demonstrated that raw motion signals, coupled with interpretable dynamical features, can meaningfully stratify sperm motility into coherent phenotypes. By integrating DTW-based clustering (73) with phase-space geometry (60, 64) and fatigue metrics (70) we revealed hidden substructures in sperm movement that reflect distinct dynamical signatures.

The resulting phenotypes were robust, biologically plausible, and interpretable in terms of progression, variability, and endurance. These findings underscore the potential of combining time-series modeling with biophysically informed descriptors to uncover latent motility profiles. Such approaches may support the development of explainable tools for sperm analysis, independent of morphology or concentration, and contribute to a deeper understanding of motility heterogeneity in the male reproductive system.

## Notes

https://zenodo.org/records/7293726

### Competing Interest Statement

The authors have declared no competing interest.

